# Pollinator habitat plantings benefit wild, native bees, but do not necessarily favor rare species

**DOI:** 10.1101/2021.05.24.445524

**Authors:** Daniel P. Cariveau, Michael Roswell, Tina Harrison, Mark Genung, Jason Gibbs, Rachael Winfree

## Abstract

1. Installing pollinator habitat is a ubiquitous conservation tool, but little is known about which pollinator taxa require support, or which benefit from habitat installations.
2. We studied the response of rare and common bees to pollinator habitat enhancement. We used independent regional datasets to designate bee species as common or rare based on their rank according to one of three metrics: a) site occurrence frequency, b) local relative abundance, and c) geographic range size. We asked whether the abundance or richness of rare and common bees were greater in pollinator habitat, relative to old-field controls. Because we used an arbitrary, quantile-based cutoff to categorize species rarity, we conducted sensitivity analysis and controlled for rarity classification errors with a null model. With this null model, we determined whether rare and/or common species responded to pollinator habitat disproportionately, compared to the expectation for ‘typical’ bee species.
3. We found that the number of individuals and of species designated as rare based on local relative abundance was greater in pollinator habitat enhancements. The number of individuals from bee species designated as rare based on site occurrence was lower in pollinator habitat enhancements, but the number of species was not clearly different between habitat types. We did not find a clear positive nor negative effect of habitat enhancement for species designated rare based on geographic range size. For all three rarity metrics, common species increased in abundance and richness in pollinator habitat relative to controls. Null models indicated that in most cases, neither rare nor common species disproportionately benefited from pollinator habitat.
4. *Synthesis and Applications:* Our results suggest that pollinator habitat can lead to an increase in the abundance and richness of bees, including species that are rare and that are common. However, rare species appeared to respond differently than typical species, and depending on how species were classified as rare, could display muted or even negative responses to habitat enhancement. Targeting rare species with specific floral resources or unique habitat types may lead to better outcomes for rare and threatened species.

## 1 INTRODUCTION

Habitat restoration is increasingly an important tool for conserving biodiversity (Hobbs 2007; Brudvig 2017). Currently, considerable government funding goes into restoring habitat for pollinators (Garbach & Long 2017). The primary method by which land managers restore habitat is the planting of wildflowers to provide resources such as pollen and nectar (Tonietto & Larkin 2018). For wild, native bee communities, the lack of floral resources is likely one of the main limiting factors (Roulston & Goodell 2011) and has been linked to declines in native bee health and abundance as well as species extinction (Carvell *et al*. 2006; Williams *et al*. 2012; Ollerton *et al*. 2014; Crone & Williams 2016). Increasing floral cover and diversity increases the overall diversity and abundance of bee pollinators (Blaauw & Isaacs 2014; Kleijn *et al*. 2015; Scheper *et al*. 2015; Williams *et al*. 2015; Tonietto & Larkin 2018; Albrecht *et al*. 2020).

Rare species comprise the bulk of biodiversity (McGill *et al*. 2007), and are often at greater risk of extirpation (Lande 1988; Manne *et al*. 1999; Payne & Finnegan 2007; Harnik *et al*. 2012). Conservation practices are often implemented to protect or benefit rare species *per se* (Hallett *et al*. 2013). However, few studies have specifically examined whether particular groups of pollinator species benefit from pollinator plantings (but see Pywell *et al*. 2012; Scheper *et al*. 2015; Sutter *et al*. 2017). There are reasons to expect that pollinator plantings both would, and would not, benefit rare species. On the one hand, such plantings could have a disproportionately positive effect on rare species if they contained specific resources that are limiting to those rare species in the wider landscape (Gibbs *et al*. 2008; Swarts & Dixon 2009). On the other hand, pollinator plantings in a degraded landscape might benefit only species that persist in these landscapes, and are thus resilient to anthropogenic change and likely not of greatest conservation concern (Kleijn *et al*. 2015). Lastly, restored pollinator plantings may similarly benefit both rare and common species. In this case, changing plant seed mixes to increase flower species preferred by rare bees may lead to better conservation outcomes for all species (Sutter *et al*. 2017; MacLeod *et al*. 2020).

An important first step in assessing the impact of habitats on rare species is to define which species are, in fact, rare. Rarity can be defined using multiple metrics, including local abundance, site occurrence frequency across a larger area, and/or geographic range size (Rabinowitz 1981; Gaston 1994). Species categorized as rare based on different metrics may be sensitive to different ecological processes. Species with low local abundance are prone to local extinction via stochastic demographic processes (Lande 1988). Site occurrence frequency can predict extinction risk (Payne & Finnegan 2007; Harnik *et al*. 2012) because species that are found at fewer sites are less likely to recolonize following disturbance (Fagan *et al*. 2002).

Further, species that are only at a few sites may be habitat specialists and at risk if those particular habitats are lost (Rabinowitz 1981). A third metric, geographic range size, is an important predictor of extinction in the fossil record (Kiessling & Aberhan 2007; Payne *et al*. 2011). Species with small ranges may be especially worthy of conservation action because populations may be less able to escape more local- or regional-scale disturbances such land-use change (Manne *et al*. 1999; Manne & Pimm 2001).

Here, we test whether pollinator habitat installations, created through wildflower plantings developed for pollinators (herein pollinator planting) generally, benefit rare and/or common species of native bees. We sampled native bees in a paired study design, in which each pollinator planting ‘treatment’ was compared to a nearby semi-natural old field ‘control’. Old fields provide a realistic control habitat type because in our study region, they are the early successional habitat that regenerates without any intervention. They also provide a conservative control for assessing the effect of pollinator plantings, because they are one of the best habitats for wild bees in the region of this study (Mandelik *et al*. 2012). To compare bee communities between pollinator plantings and controls, we collected all species of wild, native bees at 16 pairs of sites over five years. Because there are no comprehensive assessments (such as Red Lists) of bee rarity for North America, we categorized species as regionally rare or common based on large, independent datasets. We asked two questions: 1) Do pollinator habitat plantings benefit rare and/or common bees? 2) Do pollinator habitat plantings benefits accrue disproportionately to either rare or common bee species? We answered these questions categorizing species as rare or common based on one of three metrics: a) local relative abundance, b) site occurrence frequency, and c) geographic range size

## 2 METHODS

### 2.1 Study Design and Data Collection

This study took place from 2011 to 2015 in New Jersey and eastern Pennsylvania, USA (Fig. 1). We selected a total of 16 pollinator plantings that were separated by at least 4.5km. All habitats were installed by private landowners following United States Department of Agriculture National Resource Conservation Service guidelines for pollinator habitat. We used a paired design and at each pollinator planting chose a nearby old field as a control plot. Old-field control plots were located between 200-800m from the pollinator planting, which allowed us to minimize the spatial variation in native bee communities and landscape context between the habitat and controls. Hereafter, we refer to a pollinator planting and its paired old-field control together as a site.

**Figure 1.**
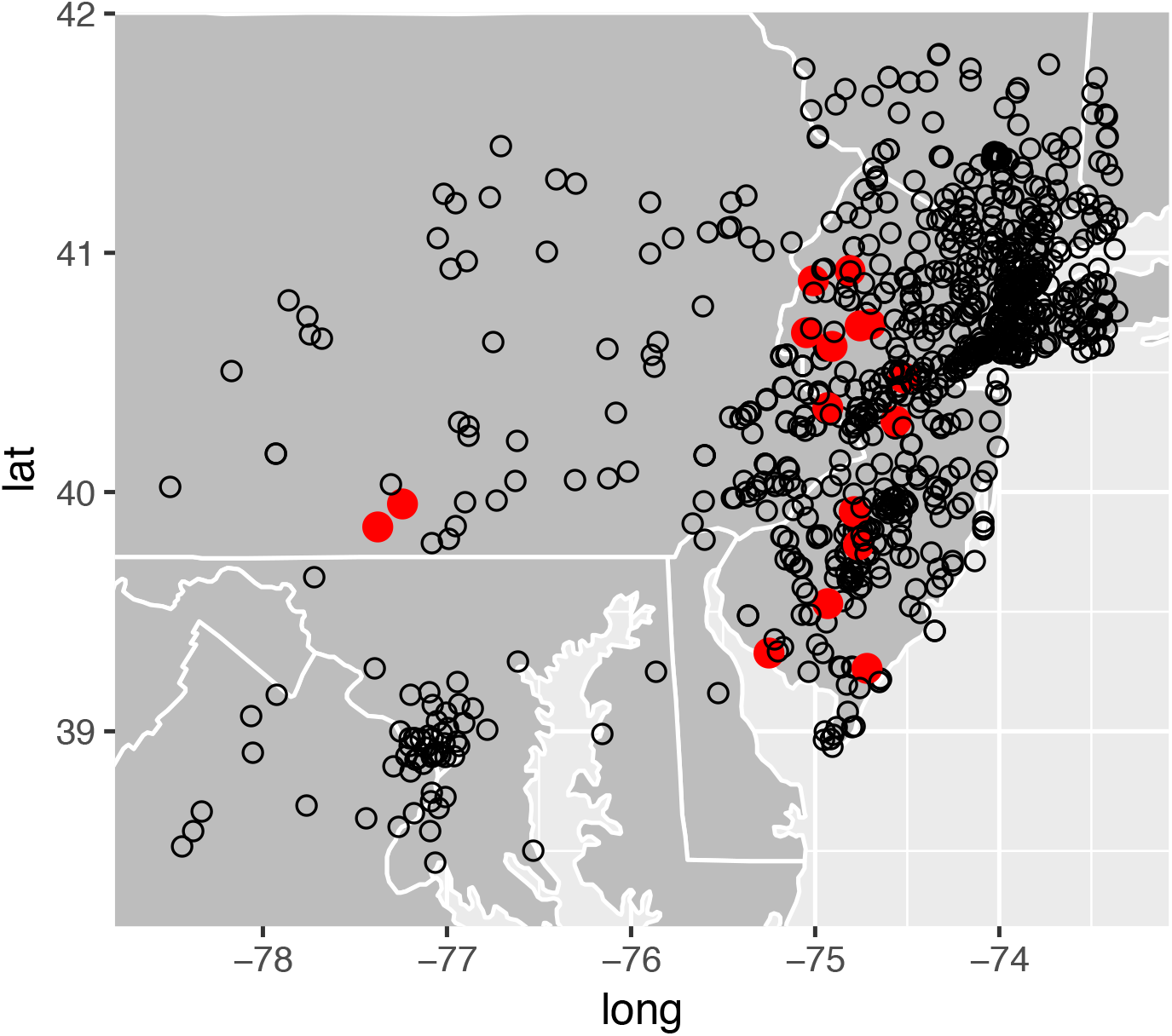
Map of wildflower plantings (solid red circles) and locations of specimen records from independent regional dataset (open black circles).

To determine bee use of pollinator planting and old-field controls, we sampled bees along 40 m transects. In any given site-by-year combination, we sampled either 2 or 4 transects, always with the same number of transects and equal effort at both the treatment and control plots within a site. We walked along each transect and collected all wild, native bees visiting flowers within 1m of each side of the transect. Transects were sampled for 10 minutes each, with the timer stopped to process specimens. Each transect was sampled once in the morning and once in the afternoon. To minimize variation due to weather, we sampled bees in dry, warm (>14C), and still conditions (winds below 3.5 m/s). To minimize observer bias, each collector completed the same number of transects in the pollinator restoration and old-field control each day. Sites were sampled approximately once a month from June through September for a total of four rounds per year. Sites were sampled for 1-4 years. This resulted in a total of over 293 hours of net collecting, not including time to process specimens.

All bees were pinned, labeled, and identified to species or species complex (Table S1). To identify bees to species, we used the following taxonomic revisions (Mitchell 1960; LaBerge 1961; Mitchell 1962; LaBerge 1967; Ribble 1968; LaBerge 1971, 1973; LaBerge and Ribble 1975; LaBerge 1977; Bouseman and LaBerge 1979; LaBerge 1980, 1986; McGinley 1986; LaBerge 1987; Laverty and Harder 1988; LaBerge 1989; Coelho 2004; Gibbs 2011; Rehan and Sheffield 2011; Gibbs et al. 2013) and keys available online (Arduser 2016; Larkin et al. 2016). With the exception of the three most common bumble bee species (*Bombus impatiens, B. bimaculatus*, and *B. griseocollis*), and the carpenter bee *Xylocopa virginica*, for which only voucher specimens were kept, all specimens are stored in the Winfree Lab at Rutgers University in New Brunswick, NJ USA. This dataset is hereafter referred to as the pollinator planting dataset and is distinct from the independent datasets used for determining rarity, described below.

### 2.2 Determining Rarity

Using independent datasets, we categorized species as rare or common based on three metrics: local relative abundance, site occurrence frequency, and geographic range size. Species were rare or common depending on whether they were in the lowest or highest quartile (Gaston 1994). Descriptions of the datasets and which datasets were used for each rarity metric follow below.

#### 2.2.1 Local Relative Abundance

To determine whether a species was rare or common in terms of local relative abundance and site occurrence, we compiled a large dataset of bee specimens collected in the states of New Jersey, Pennsylvania, and New York, USA. The specimens came from six studies, conducted by members of the Winfree research group, in which bees were collected using pan traps or hand nets between 2003 and 2016 (Table S2). In all of these studies, collectors did not target particular species or species groups, and collection took place irrespective of abundance; therefore, we assumed that specimen records from these datasets, on average, reflect actual bee abundances at sites in our region, and use these records to compute relative local abundance within sites.

We filtered records to generate rarity designations for our study region using the following criteria. First, to evaluate rarity within the same region as the pollinator plantings, we drew a rectangle where each edge was 100km from the nearest site in each cardinal direction. Second, we filtered records for phenology, and used only records collected within two weeks of the year from the start and end of the collection events of our restoration study. This could mean that in our filtered dataset, bee species that are common either outside of our spatial scale or in a different season may be classified as rare. Third, we removed exotic species (1.4% of all individual records and <5% of species across the 6 reference data sets). Our study focused exclusively on native species for a number of reasons. First, both the abundance and site occurrence frequencies of exotic species are likely driven by unique factors such as human-mediated colonization events and time since invasion. Second, as many of our specimens from the independent datasets come from historic collections, exotic species are likely underrepresented. Finally, we were most interested in characterizing native species; exotic species are rarely the target of conservation efforts.

We then calculated local relative abundance as follows: First, to avoid artifacts related to small sample sizes, we removed all records from sites at which fewer than 10 total specimens had been collected. Second, we calculated relative abundance for each species at each location. Abundances for each species at each locality were summed across years and sample rounds. Third, for each species, we calculated mean local relative abundance across sites by summing the relative abundances at all sites at which a species was present, and dividing this sum by the number of sites at which that species was detected. Thus, when a species was not recorded at a site, that site was not included in the local relative abundance calculation for that species; this exclusion of zeros was done to make our measure of relative abundance distinct from site occupancy (below). Finally, there were 10 species (represented by 47 specimens) that were collected in the pollinator plantings dataset but were not recorded at the sites used to calculate relative abundance. These species were removed from the analysis for local relative abundance. In total, the data used to calculate local abundance consisted of 52,885 specimens and 294 species. Collections took place at 261 different locations (Fig. 1).

#### 2.2.2 Site Occurrence Frequency

To assess rarity based on site occurrence frequency, we used the same dataset used for local relative abundance (see above), supplemented by a previously published dataset compiled from species records from nine museums (Bartomeus *et al*. 2013). This dataset includes over 11,000 collection events, spanning 139 years (1870-2010). The data set is filtered to use only one record (individual bee) per species per collection event to reduce biases that might be present in museum collections. Specifically, collectors were unlikely to collect each species proportional to its relative abundance. Thus, we use this data set for the occupancy analysis but not the abundance analysis (above). For use in the occurrence analysis, we standardized the abundance dataset to presence-absence form also (i.e., retained only one individual of each species per site-date). We filtered the museum dataset for phenology, spatial extent and native status in the same manner as the abundance dataset. After filtering and combining, the occurrence data set included 59,918 individual records of 384 species across 992 different locations (Fig. 1). To determine site occurrence frequency, we counted the number of sites at which each species was collected. Finally, we collected five species (represented by 35 specimens) in the pollinator habitat dataset but that were not recorded at all in the site occurrence dataset. We categorized these species as rare based on site occupancy.

#### 2.2.3 Geographic Range Size

To assess geographic range size, we used the methods from a study completed by some of the present coauthors (Harrison *et al*. 2017). This study used recorded state records from the Discover Life AMNH_BEES database (Ascher & Pickering 2019). A minimum bounding polygon was then calculated to contain the centroids of each state in which each species was present. For details see (Harrison *et al*. 2017).

### 2.3 Quantile Designation of Rare and Common Species

For *local relative abundance*, 74 species in the regional dataset were designated as rare (local relative abundance = <0.81%) and 73 species as common (local relative abundance = >3.7%). For *site occurrence frequency*, there was an unequal number of species in some quartiles due to ties. Ties for the first and second quartiles and the third and fourth quartile were placed in the first and fourth quartile respectively. There were 105 species designated as rare (< 4 sites) and 94 as common (>26 sites). For *geographic range size* we designated 96 species as rare and 96 as common. A table of all species and their rarity classifications can be found in Table S1.

### 2.4 Data Analysis

#### 2.4.1 Do pollinator habitat restorations benefit rare and or common bees?

We designated each species collected in the pollinator restoration dataset as rare or common (or neither, constituting those between the 25th-75th percentiles) for each of the three rarity metrics. For each pollinator planting and old-field control at each site, we then summed (across all years and sampling rounds) the number of individuals (hereafter, “abundance”) and species (hereafter, “richness”) of rare and common bees. To determine how pollinator planting affected the richness or abundance of rare, and separately, common bees, we fit generalized linear models with either richness or abundance of either rare or common bees, according to one of the three metrics, as the response variable. As predictors in each model, we incorporated treatment (pollinator planting or old-field control) as a fixed effect. Additionally, because we used a paired study design, with one wildflower planting and one control at each site, we included site as a random effect. We fit generalized linear mixed-effects models for each response variable with a negative binomial error structure using the function *glmer_nb* from the R package “lme4” (Bates *et al*. 2015). Model assumptions were checked using the DHARMa package (Hartig 2020). This analysis was conducted using R 4.0.5 (R Core Team 2021).

#### 2.4.2 Do pollinator habitat restoration benefits accrue disproportionately to either rare or common bee species?

The models described above test for increases in the abundance or richness of a specified group of species (e.g. those that have smaller geographic ranges), regardless of how other species respond. If managers believe that rare species are especially vulnerable to global change, it may be strategic to focus management actions that benefit these species. We generated a null model to test whether the species we designated as rare, or those we designated as common, especially benefit from habitat enhancement, or whether they respond about as strongly as expected, given that the response group consists of about one quarter of the species in our study region.

In each iteration of our null model, we designate a random subset of species as rare or common, where the number of rare and common species in the null group is the same as the number of rare and common species in the empirical data. We used our observations of these species in the pollinator habitats and old-field controls, and refit the models for each rarity criterion (i.e. for each group of [null] “rare” or [null] “common” species) and each response (i.e. richness or abundance). We then compared the model estimates for the effect of habitat enhancement for the species group from 9,999 iterations of each null model to the model estimate for our observed data. In short, this null model tells us how strongly any randomly-chosen fraction (up to ∼25%, depending on overlap between regional rarity and occurrence in the restoration dataset) of our species responds to habitat enhancement, providing scope to test whether rare species respond especially strongly to habitat enhancements. There are at least two reasons that we could fail to find a clear difference between the empirical model coefficients and those from the null models. The first of these is biological: the factors leading species to be regionally rare or common may not be strongly correlated to the factors that predispose species to respond strongly to habitat enhancement. The second of these is more related to classification accuracy: Even if *truly* regionally rare and common bee species respond differently to habitat enhancement than typical species do, we may have misclassified species as rare or, less likely, as common, obscuring the link between rarity/commonness and the propensity to benefit from habitat enhancement (Harrison *et al*. 2017). This null model analysis entailed generating nearly 120,000 mixed-effects models, and therefore, we did not complete model validation for the null datasets.

We consider a group to clearly exhibit a stronger or weaker than expected response to pollinator habitat restoration (e.g., abundance in pollinator planting vs. abundance in control) when empirical model estimates are above 97.5% or below 2.5% of the null model results. This analysis was conducted using R 3.4.1 (R Core Team 2017).

#### 2.4.3 Testing sensitivity of quartiles

To test whether our results were sensitive to the quantile threshold for what constitutes rarity (or commonness), we re-ran our analyses and the null model for 10 additional thresholds, ranging from the 20th to the 30th percentile. This analysis was conducted using R 3.4.1 (R Core Team 2017).

## 3 Results

In the pollinator restoration dataset, we collected a total of 10,809 individual bees of 157 native species or species complexes. Of these, 195 specimens (37 species) were designated as locally rare and 8,642 specimens (39 species) as locally common, based on the relative abundance data set. Based on the site occurrence dataset, 65 specimens (14 species) were designated as rare and 9,977 specimens of (77 species) were designated as common. For the geographic range size metric, 316 specimens (34 species) were rare and 4081 specimens (40 species) were common.

### 3.1 Do pollinator habitat restorations benefit rare and/ or common bees?

Species defined as rare based on local relative abundance had higher abundance and richness? in the pollinator plantings compared to old-field controls by factors of nearly 3 and 2 (Table 1; Fig. 2A). Species categorized as rare based on low site occurrence frequency did not exhibit a clear difference in the number species collected; however, the abundance of these rare bees was reduced by 50% reduction in treatment versus controls (Table 1, Fig. 2B). Species categorized as rare based on small geographic range size were not clearly more abundant or species rich in pollinator plantings (Table 1, Fig. 2C). Regardless as to how we defined commonness, we collected both more species and individuals from common bee species in pollinator restorations compared to controls (Table 1, Fig. 3).

**Table 1.**
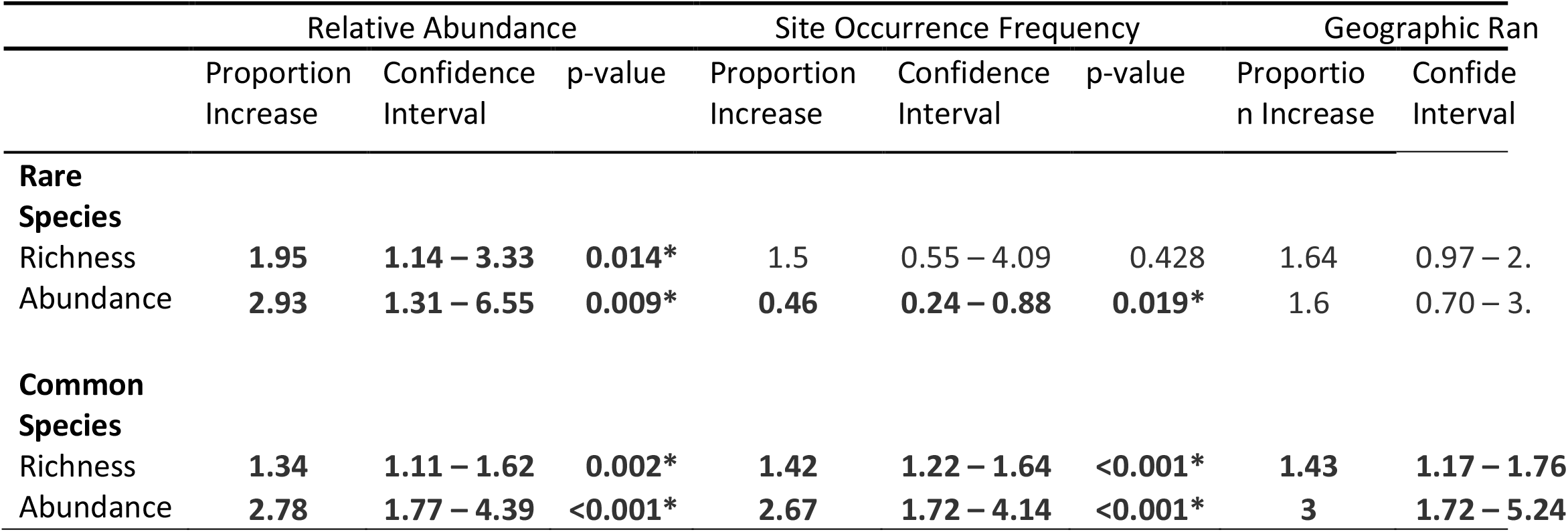
Model output of comparison between pollinator plantings and paired old-field controls for all three rarity metrics. Proportion change is exponentiated model output of change in pollinator planting treatments compared to old-field controls. Bold and asterisks represent significant difference between pollinator planting and old filed controls at p < 0.05.

**Figure 2.**
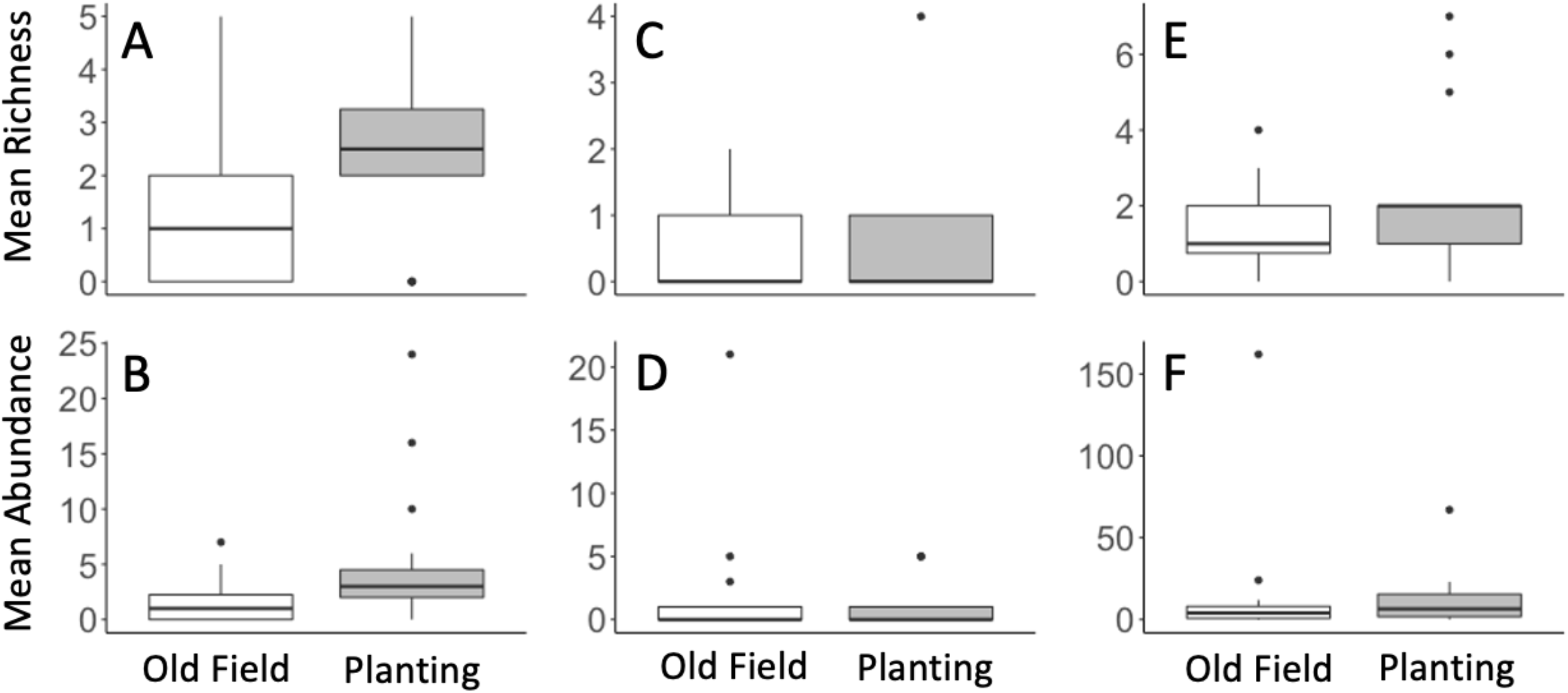
Rare species richness and abundance in old fields (white) and pollinator habitat (grey). Rarity metrics for A-B) Relative abundance, C-D) site occurrence frequency, and E-F) geographic range size.

**Figure 3.**
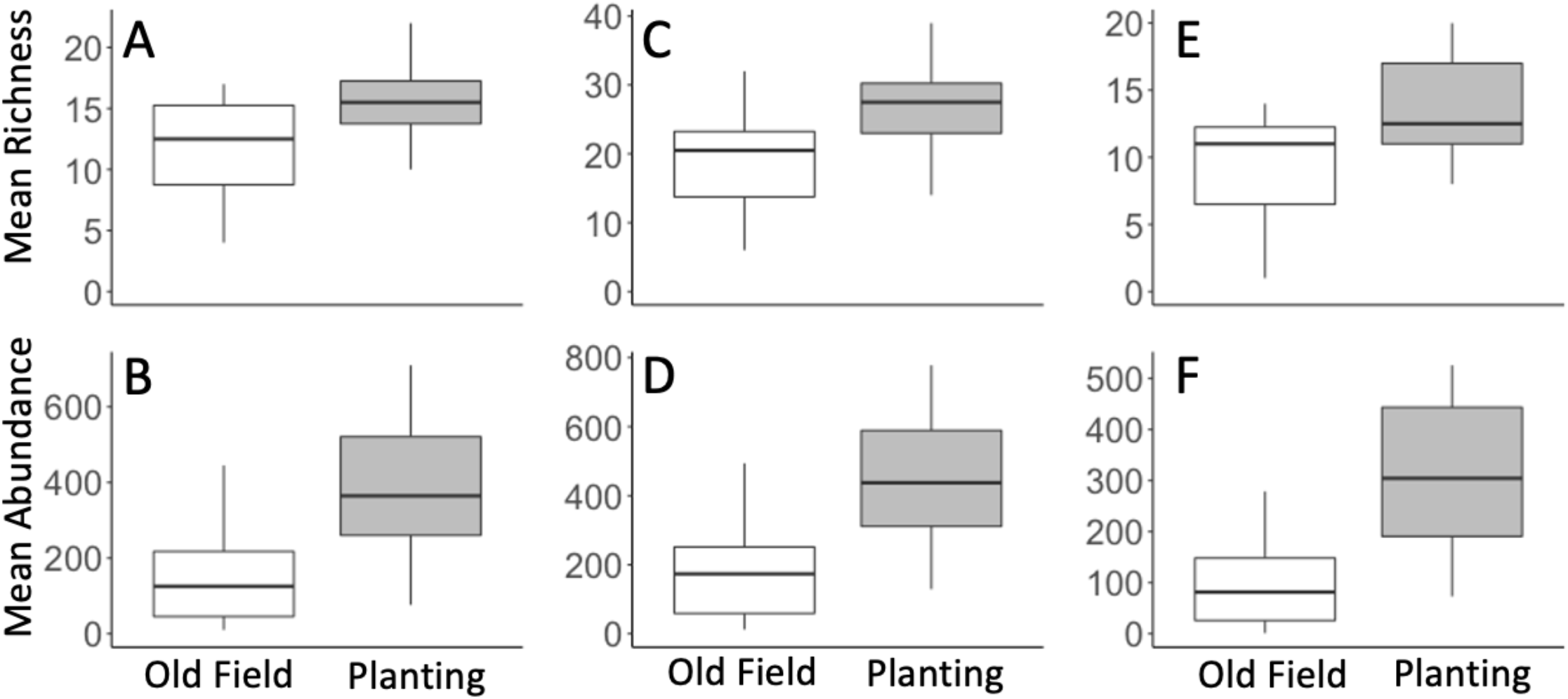
Common species richness and abundance in old fields (white) and pollinator habitat (grey). Rarity metrics for A-B) Relative abundance, C-D) site occurrence frequency, and E-F) geographic range size.

### 3.2 Do pollinator habitat restorations lead to a disproportionate benefit for either the rare or the common bees?

For almost all species rarity criteria and response types (richness or abundance), we found no difference between the response of the species designated as rare, and results for a null model that randomly chose an equally-sized group of species (Fig. 4). There was one exception: habitat enhancements supported 54% lower abundance of bee species designated as rare based on regional site occupancy (Fig. 4e, p<0.01). Similarly, for bee species designated as common (for all commonness criteria) richness and abundance responses to habitat enhancement were not different than expected for a random group of species (Fig. 5).

**Figure 4.**
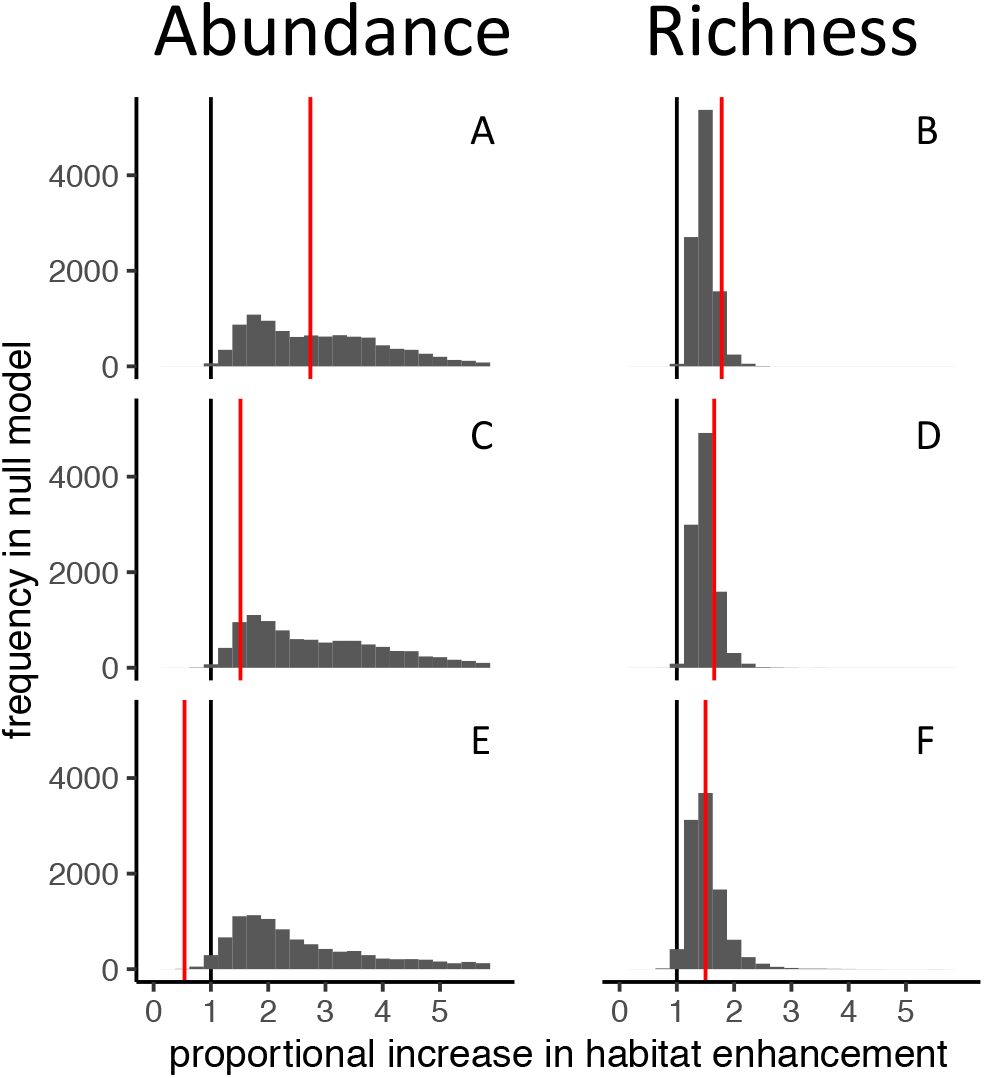
Comparison of outputs for model simulations and empirical models for rare species. x-axis values represent proportional increase in abundance or richness in habitat enhancements relative to old field controls. Histograms represent the frequency of each proportional change in richness or abundance in the null model, which designates randomly selected species as “rare.” Black lines represent no difference in response variable between treatment and control (x=1). Red lines represent model output for the real data, for which species were defined as rare based on regional relative abundance (A,B), regional site occurrence frequency (C,D), or geographic range size (E, F).

**Figure 5.**
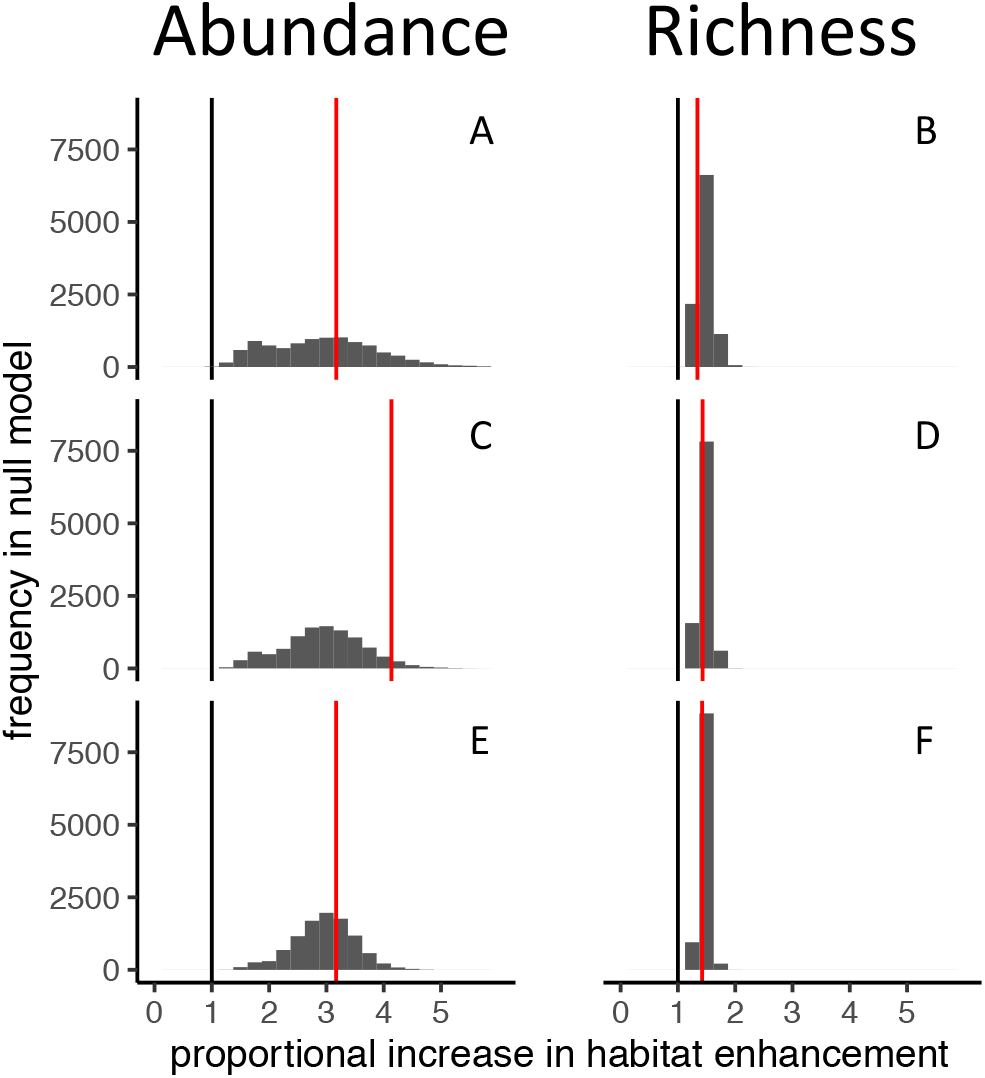
Comparison of outputs for model simulations and empirical models for common species. x-axis values represent proportional increase in abundance or richness in habitat enhancements relative to old field controls. Histograms represent the frequency of each proportional change in richness or abundance in the null model, which designates randomly selected species as “common.” Black lines represent no difference in response variable between treatment and control (x=1). Red lines represent model output for the real data, for which species were defined as common based on regional relative abundance (A,B), regional site occurrence frequency (C,D), or geographic range size (E, F).

### 3.3 Testing sensitivity of quartiles

The quantile sensitivity analysis mostly did not reveal qualitative differences in the results (Table S3). However, as the quantile thresholds for rarity/commonness got larger (and thus more inclusive), model coefficients tended towards the average for all species.

## 4 Discussion

### 4.1 Rare bee species exhibit atypical responses to habitat enhancements

Pollinator plantings, in general, increase the overall abundance and richness of wild, native bees (reviewed in Tonietto & Larkin 2018; & Nicholson *et al*. 2020). We found that common bee species and, depending on the criteria for designation, rare bee species had higher abundance and species richness in wildflower plantings as compared to old-field controls. Our results partially support other studies that found pollinator plantings benefit wild bees, including rare and common species (Pywell *et al*. 2012; Scheper *et al*. 2015). For example, in the EU, Scheper et al. (2015) found that all bee species, including those designated as rare by the IUCN, increased in floral strips. Pollinator plantings likely benefit both rare and common species when both groups overlap in the species of flowers they prefer, and these floral species are included in the plantings (Macleod *et al*. 2020). As common species are among the most important crop pollinators (Winfree *et al*. 2015), pollinator plantings that benefit them (even if they do not also support rare species effectively) could have positive impacts on crop pollination (Blaauw & Isaacs 2014; but see Nicholson *et al*. 2020; Lowe *et al*. 2021). However, depending on how bee species were categorized as rare, and which response variable we tracked, pollinator plantings failed to benefit all rare bees. In particular, bee species categorized as rare based on regional site occurrence frequency had lower abundance in pollinator plantings as compared to old-field controls. Overall, this indicates that, although pollinator plantings in general can have benefits for biodiversity and ecosystem services, these two goals are not always met simultaneously (Nicholson *et al*. 2020; Lowe *et al*. 2021), and not all species benefit from this conservation action (Senapathi *et al*. 2015).

The effect of pollinator plantings on rare species depended on rarity criteria. Bee species designated as rare based on local relative abundance were more abundant and species rich in pollinator plantings. Even if a species has low local abundance, it may still benefit from pollinator plantings if it occurs frequently across the landscape. This echoes findings elsewhere that benefits of pollinator habitat enhancement are greatest in landscapes regions with intermediate cover of natural and seminatural habitat, and are therefore likely to contain source populations able to exploit additional resources (Grab *et al*. 2020; Griffin *et al*. 2021; McCullough *et al*. 2021, but see Lane et al. 2020).

Conversely, species that do not commonly occur across a region may be unlikely to benefit from pollinator plantings for two reasons. First, a number of rare bees are specialists and collect pollen from a limited suite of plants. These species may be common when their host species is present. Wildflower seed mixes used in pollinator plantings are only a small subset of native plants within a region, inevitably fail to include important floral hosts. For example, at one site, we caught seven *Trachusa dorsalis* (Megachilidae; Lepeletier 1841) in an old-field control. This species had not been recorded in New Jersey or Pennsylvania (states where this study occurred) for over 100 years (Ascher & Pickering 2019). *Trachusa dorsalis* specializes on plants in the genus *Strophostyles* (Fabaceae), a genus not planted in the pollinator habitats we sampled. Second, plantings are not targeted towards specific habitat types that may support rare bees (Harrison *et al*. 2018). In our dataset, specialist bee species are likely to be categorized as rare based on site occurrence frequency (assuming the resource on which they specialize is heterogeneously available across the region), and potentially by approximate range size (if the resource on which they specialize is also geographically restricted). We found that bee species with low regional site occupancy showed much less of an abundance response to habitat installation than we would expect for a random group of species, in line with our expectation that pollinator plantings may not benefit uncommon specialist species with atypical resource requirements. This finding echoes suggestions elsewhere (e.g. Macleod *et al*. 2020) that species-specific management actions may be necessary if rare species conservation is an objective of restoration efforts.

### 4.2 Limitations

One limitation of our study is that the plantings we studied did not contain early-blooming spring plants - a common problem in pollinator plantings (Wood et al. 2018). Therefore, we were not able to assess whether pollinator plantings with spring plant species might benefit spring bees. This may be particularly important in our region, as a number of early spring species are known to be rare (Harrison *et al*. 2017).

A more general limitation is the inherent problems with categorizing species as rare in the first place. Here we use a number of metrics and employ a quantile method to characterize species rarity (Gaston 1994). This method has a number of important advantages: it is straightforward, has a clear cut-off, and easily incorporates new data. One potential drawback is the need to make an arbitrary decision about which quantile value to use for rarity designation (Gaston 1994). Here we show that our results were robust to choice of quantile. However, there are other important considerations when using this approach. For example, sampling biases (e.g. in habitat type) in the reference datasets means the quantitative rankings might still not reflect wider patterns of abundance and occurrence in nature.

We faced both misclassification risks and statistical limitations inherent to studying rare species. Due to limited monitoring efforts for wild bees, it is not clear which species in our region are truly rare, and classification errors are likely (Harrison *et al*. 2017). First, in defining rarity, we chose to focus on records collected within a few hundred kilometers of our study sites, and within a few weeks of the year of our sampling period, and thus consider species to be rare even if they are common elsewhere or in other seasons. Second, due to limited data coverage, it is likely that our rarity categorizations conflated rarity types: species with low abundance (for example, specialized kleptoparasitic bees) could be mistakenly categorized as low occupancy even if they are widespread in our study region, or species with strong habitat specialization or low relative abundance over a large range could have erroneously small range-size estimates. Furthermore, binary rarity classifications based on arbitrary thresholds (and for regional metrics, arbitrary spatial scale) are likely to miss more biologically relevant definitions of rarity (Gaston 1994; Inger *et al*. 2015; Reed *et al*. 2020; Balbuena *et al*. 2021). Additionally, because they are rare, any study of rare species is likely to face issues of statistical power and related concerns, such as type-S and type-M errors (Gelman & Carlin 2014). In part, we were able to defend against these issues with sensitivity analysis and our null model simulation. Yet, even as qualitative patterns were robust to changing rarity thresholds, and null-model simulations, in some cases, indicated that differing responses from rare species were not expected simply as a result of classifying some species as rare and others as not rare, we urge caution in generalizing our findings. Overall, our results demonstrate heterogeneity in response to habitat enhancements at the species level, and many species are, of course, rare. Diverse taxonomic and functional groups are unlikely to respond uniformly to conservation actions, and more targeted conservation goals and strategies may be easier to assess and fine-tune.

### 4.3 Conclusions and Management Implications

Currently, plant species used in pollinator plantings are chosen because of cost, availability, or whether the seed mix attracts high pollinator species richness or abundance (Harmon-Threatt & Hendrix 2015; M’Gonigle *et al*. 2017; Williams & Lonsdorf 2018). However, it may be that current pollinator plantings primarily benefit species that already persist within anthropogenic landscapes (Senapathi *et al*. 2015). If land managers aim to conserve rare bees, they may need to incorporate plants not traditionally used in pollinator habitat plantings (MacLeod *et al*. 2020).

Although they were the focus of our study, it is not necessarily the case that rare species *should be* the target when designing and implementing pollinator plantings. While conservation biology has focused on rarity, temporal trends (population decline, habitat loss, and range contraction) may better predict extinction risk (Collen *et al*. 2016) – of course, these trends are difficult to detect in rarely sampled species, such as many we have studied here. At the same time, common species are also of critical conservation concern. In Europe, common bird species are experiencing declines while less abundant species are increasing (Inger *et al*. 2015). Further, common species may also have large effects on ecosystem function (Gaston 2011; Kleijn *et al*. 2015; Genung *et al*. 2018). More important than whether current practices benefit rare species, ensuring overlap between conservation approaches and their likely impact on *particular* species of concern may produce the best conservation outcomes, as disparate responses to habitat alterations should be expected across diverse taxonomic groups.

## Author Contributions

RW, MR, and DPC designed the study. DPC, MR, MG, and TH collected the data. DPC wrote the manuscript with considerable input from MR. All authors contributed to writing and conceptual development. MR and DPC analyzed the data. RW oversaw writing and analysis. JG identified native bee specimens.

## Acknowledgements

Colleen Smith and Molly MacLeod contributed data analysed in this study. We thank Rosy Tucker, Kurtis Himmler, Joe Zientek, Elena Suglia, Abigail Cohen, Gabriella Pardee, Stevia Morawski, Chava Weitzman, Jeff Bogdewitz, Bridget Johnson, and Geetha Nayak for their assistance with data collection, and the landowners who supported this work.

## Table S1. Supplementary Information on the datasets used to classify rarity

### Dataset Name: dryad_amnh

Use in manuscript: site occurrence frequency

Number bee individual specimens used in this study: 7633

Number bee species used in this study: 331

Number of sites: 751

Published in: (Bartomeus *et al*. 2011, 2013a, b)

### Dataset Name: bef_scale_spec

Use in manuscript: site occurrence frequency, local relative abundance

Number bee individual specimens used in this study: 7359

Number bee species used in this study: 90

Number of sites: 25

Unpublished

### Dataset Name: cape_may

Use in manuscript: site occurrence frequency, local relative abundance

Number bee individual specimens used in this study: 8217

Number bee species used in this study: 85

Number of sites: 1

Published in: (MacLeod *et al*. 2016, 2020; Genung *et al*. 2017)

### Dataset Name: forest_spec

Use in manuscript: site occurrence frequency, local relative abundance

Number bee individual specimens used in this study: 543

Number bee species used in this study: 59

Number of sites: 33

Published in: (Smith *et al*. In Press.; Volenec & Smith 2021)

### Dataset Name: male_bee_project

Use in manuscript: site occurrence frequency, local relative abundance

Number bee individual specimens used in this study: 15110

Number bee species used in this study: 145

Number of sites: 8

Published in: (Roswell *et al*. 2019a, b)

### Dataset Name: njpa_ha

Use in manuscript: site occurrence frequency, local relative abundance

Number bee individual specimens used in this study: 4822

Number bee species used in this study: 72

Number of sites: 54

Unpublished

### Dataset Name: nsf0607_spec

Use in manuscript: site occurrence frequency, local relative abundance

Number bee individual specimens used in this study: 683

Number bee species used in this study: 70

Number of sites: 18

Published in: (Winfree *et al*. 2014)

### Dataset Name: pinelands_2003

Use in manuscript: site occurrence frequency, local relative abundance

Number bee individual specimens used in this study: 1967

Number bee species used in this study: 124

Number of sites: 44

Published in: (Winfree *et al*. 2007)

### Dataset Name: bh_spec

Use in manuscript: site occurrence frequency, local relative abundance

Number bee individual specimens used in this study: 5641

Number bee species used in this study: 162

Number of sites: 27

Published in: (Harrison *et al*. 2017, 2018b, a)

### Dataset Name: swg

Use in manuscript: site occurrence frequency, local relative abundance

Number bee individual specimens used in this study: 7943

Number bee species used in this study: 192

Number of sites: 51

Unpublished

